# Assessing Enclosure Modifications to Enhance Accessibility for Zoo-Housed Meerkats (*Suricata suricatta*)

**DOI:** 10.64898/2026.11.04.764381

**Authors:** Alehna Crowe, Ethan Mead, Shannon L. O’Brien

## Abstract

Scientific advancements have contributed to increased longevity of animals in zoos and aquariums, creating a growing need for environments that remain accessible to geriatric animals and animals with disabilities. However, few empirical studies have evaluated whether accessibility-focused enclosure modifications impact species-appropriate behavior. We evaluated the effects of accessibility modifications on a group of four adult zoo-housed meerkats (*Suricata suricatta*), including one individual with a forelimb amputation and one geriatric individual. Low-level logs and steps were added to the enclosure to provide intermediate heights and facilitate access to elevated areas. Following the introduction of these enclosure modifications, sentry behavior increased, with all individuals engaging in this behavior compared with only one individual during baseline. The meerkats also increased their overall use of low-elevated areas following modification, although individual responses varied. Posture differed between study phases, with meerkats more frequently adopting tri/quadrupedal postures following modification. Together, these findings suggest that relatively simple accessibility-focused modifications can expand opportunities for geriatric animals and animals with disabilities to engage in species-appropriate behavior. More broadly, this case study highlights the importance of empirically evaluating accessibility interventions and incorporating universal design principles into animal spaces to accommodate individuals with diverse physical abilities and promote positive animal welfare.

## Introduction

Scientific advancements in fields such as veterinary medicine (Chapman and Dobbs 2023), nutrition (Cabana and Plowman 2023), and animal welfare (Binding et al. 2020) have positively contributed to the longevity of animals in accredited zoos and aquariums (hereafter, zoos) well beyond what is typically experienced by their conspecifics in the wild (Krebs et al. 2018). Similarly, these advancements have contributed to many zoos caring for animals with disabilities, defined here as the loss of a critical adaptation that has impacts on an animal’s mobility, stability, typical behavior, or perception of their environment (O’Brien 2026). A recent survey of institutions accredited by the Association of Zoos and Aquariums (AZA) found that 98% of institutions surveyed had at least one animal with a disability in their care (O’Brien 2026). Physical, cognitive, and/or sensory changes associated with advanced age or disability may impact welfare for animals in managed care if these individuals are not able to fully engage in species-appropriate behaviors (O’Brien 2026; Krebs et al. 2018). For example, animal welfare may be compromised if an individual has difficulties navigating their environment, accessing resources and/or engaging with conspecifics.

Accordingly, designing—or minimally retrofitting—animal spaces to enhance accessibility for animals who are geriatric or living with a disability can promote positive welfare (O’Brien 2026; Coe 2023). Universal design principles (Story 2001), while typically applied to human-centered spaces, can be applied to environments used by animals in managed care to promote species-appropriate behaviors. Principles of universal design include designing with accessibility in mind so that environments can be comfortably and efficiently used by individuals with diverse abilities (Story 2001). Examples of accessible design in animal spaces may include ramps or low-graded steps to elevated areas, incorporation of resting areas along movement pathways, and variation in substrate texture to enhance grip and reduce slipperiness (O’Brien 2026). Well-designed animal spaces may help prevent or delay the onset of certain age-related disabilities (e.g., limited mobility) while providing animals with continued opportunities to express choice and control (Brando and Chapman 2023).

While several empirical studies have demonstrated that enclosure design can influence species-appropriate behavior in zoo animals (e.g., de Azevedo et al. 2023; Spain et al. 2020; Earl et al. 2020), almost no published studies have directly evaluated whether accessibility-focused modifications designed for geriatric animals or animals with disabilities encourage species-appropriate behaviors or promote positive welfare outcomes (although see Zucker et al. 1991). Examining the effectiveness of enclosure accessibility modifications will advance efforts to provide animals with evidence-based care (Rose 2025) and support the development of best-practice guidelines for managing geriatric animals and animals with disabilities. Evaluating whether accessibility modifications provide opportunities to engage in species-appropriate behaviors provides a means of assessing their potential welfare benefits. Sentinel behavior in meerkats—an important vigilance behavior in which a meerkat scans the environment for predators or other potential threats while their groupmates pursue other activities such as foraging (Manser et al. 2002; Manser 1999)—offers a useful model for examining this relationship because the behavior often occurs from an elevated position (e.g., on top of rocks or logs) to maximize the sentinel meerkat’s viewing capabilities (Tatalovic 2010; Moran 1984). As such, meerkats with limited mobility, either due to aging or a disability, may have difficulty expressing this behavior.

Despite the absence of predators in managed care, meerkats in zoos have been documented performing sentinel behavior at similar rates as their wild conspecifics (Huels and Stoeger 2022). Sentinel behavior, while primarily an anti-predator strategy, also supports group cooperation (Rauber and Manser 2018; Santema and Clutton-Brock 2013). If individuals are unable to perform sentinel behavior due to advanced age or disability, social cohesion within the group may be impacted. Meerkat enclosures in zoos that allow all individuals—including those with a disability or of advanced age—to engage in sentinel behavior are likely to support positive animal welfare.

In this study, the behavior of four adult meerkats (*Suricata suricatta*; one male and three females) was evaluated before and after enclosure modifications intended to provide access to elevated areas and support sentinel behavior. One individual in this group, Alanis, had a physical disability resulting from surgical amputation of her left forelimb. In addition, all individuals in this group were at or near the median life expectancy of meerkats in zoos (9.6 years, Che-Castaldo et al. 2019). Given the advanced age of the group and Alanis’ physical disability, we predicted that the meerkats would spend more time in sentinel behavior, make greater use of elevated areas, and adopt tri/quadrupedal postures more often following accessibility-related enclosure modifications.

## Methods

### Data collection

Baseline data were collected between July 6 – 30 2025 and post-modification data were collected between July 7 – 31 2026. Collecting data during the same time of year ensured that zoo visitor activity was comparable between each phase (2025: 5 ± 4 visitors near meerkat enclosure; 2026: 6 ± 5 visitors). At the time of this study, a group of four adult meerkats was housed at Lincoln Park Zoo (Chicago, Illinois, USA) in an indoor enclosure (268 ft^2^). One female, Alanis (9 years old at study onset), underwent surgical amputation of her left forelimb following the diagnosis of a bone lesion in April 2024 (one year before study onset). A second female, Queen, was geriatric (13 years old), while the third female, Miranda (9 years old), was Alanis’s littermate. The sole male, Sonny, was 11 years old. During the baseline data collection phase, the meerkat enclosure was unmodified and included a large, elevated rock structure designed to mimic a natural termite mound (Figure 1A). The meerkats had lived in this enclosure for multiple years prior to the onset of this study. During the post-modification data collection phase, the meerkat enclosure was modified to include multiple low-level logs (Figure 1B) and steps (Figure 1C) to provide intermediate height options between the ground-level and the top of the termite mound.

**Figure 1.**
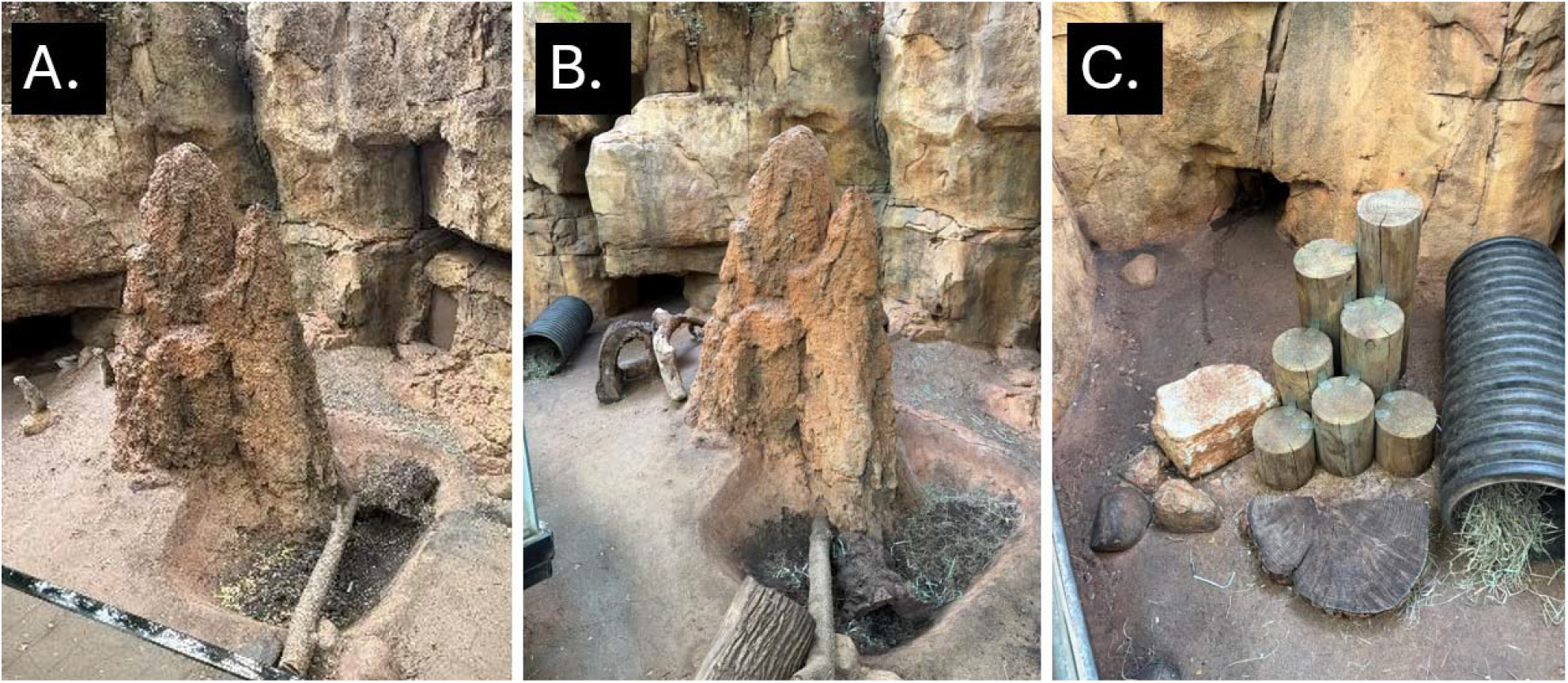
A. Meerkat enclosure at Lincoln Park Zoo (Chicago, IL, USA) during baseline data collection phase. B. Meerkat enclosure with the addition of low-level logs to provide intermediate height options for the meerkats. C. Steps and low rocks to provide additional intermediate height options.

Data collection occurred on Mondays-Thursdays between 10:00-16:00, when the building was open to the public. Each data collection observation lasted 10-minutes with data collected using interval sampling with 1-minute intervals (maximum of 10 data points collected per observation). Meerkats were observed individually in random order (10-minutes per meerkat for a maximum of 40 minutes of data collection if all meerkats were visible in the enclosure). Data collection occurred 2-4 times per day. At each 1-minute interval, meerkat behavior, height use, and location in their enclosure were recorded using the ZooMonitor app (Wark et al. 2019) on a handheld tablet device. Height use was denoted as either ground-level (not-elevated), low (slightly elevated, on low-level rocks, logs, or steps), or high (elevated on the rocky termite mound). The behavioral ethogram included 23 behaviors (Table 1) that were selected after preliminary observations of these meerkats and reviews of published studies on meerkat behavior in zoos (e.g., Bähler et al. 2024; Huels and Stoeger 2022; Sherwen et al. 2014). The behaviors standing and sitting had a follow-up question to indicate the posture of the meerkat (bipedal or tri/quadrupedal). In addition to behavioral data and height use, meerkat locations were plotted on a digital map, which converts these locations to coordinates based on a 600 x 600 coordinate point system (Wark et al. 2019). Data were collected by 6 trained observers who had previously established 100% agreement on meerkat identification and a minimum of 85% inter-observer reliability for meerkat behavior and space use within 30 coordinate points.

**Table 1.** Meerkat ethogram.

| Behavior | Description |
|---|---|
| Vocalization | Producing sounds through intentional vibration of air for the purpose of communication. |
| Sentry | Positioned on an elevated area of the enclosure and actively scanning the environment, typically sitting erect or standing, vocalizing occasionally. |
| Standing (Posture: Bipedal or Tri/Quadrupedal) | Upright with weight supported by legs and not performing another behavior listed. |
| Sitting (Posture: Bipedal or Tri/Quadrupedal) | Seated with weight supported on buttocks and not performing another behavior listed. |
| Lying | Lying with weight supported on side, back, or stomach and not performing another behavior listed. |
| Leaning | Front of body leaning on an object to support body weight, while back legs are in a sit or stand position and not performing another behavior listed. |
| Huddling | Situated in body contact of another individual and not performing another behavior listed. |
| Walking | Walking through exhibit at least a distance greater than one body’s length. |
| Running | Running through exhibit at least a distance greater than one body's length. |
| Climbing | Propelling up a vertical surface to travel a distance greater than one body's length. |
| Feeding/Foraging/Drinking | Chewing food, drinking, manipulating food for the purpose of feeding, or searching for food. |
| Digging | Using body in repetitive manner to excavate substrate (typically forelimbs but may include hind legs or snout). |
| Exploratory Behavior | Engaging with the environment, including inspecting and manipulating items. |
| Object Manipulation | Moving a non-fixed object in the enclosure using mouth or paws. (If object is associated with food, score as Feeding/Foraging/Drinking. If animal plays with object, score as Play.) |
| Play | Animal is in a relaxed state and voluntarily performing a behavior in a repetitive fashion without a clear functional benefit in the current context. |
| Self-Maintenance | Self-directed behaviors, such as stretching, yawning, rolling, or rubbing against objects (If rubbing scent glands on a surface, score as Scent Marking). |
| Scent Marking | Intentionally marking an area by rubbing scent glands on a surface. |
| Social Grooming | Licking or manipulating the skin or fur of another individual in an affiliative context. |
| Stereotypic Pacing | Animal is walking in a repetitive manner along a fixed path without an apparent goal or function. The animal must move along the path three times to qualify as pacing. |
| Reproductive Behavior | Engaged in sexual behavior with another individual. |
| Keeper Interaction | Directly interacting with keepers, either performing trained behaviors in the context of a training session or engaged in an informal manner, including feeding or inspecting. |
| Behavior Obscured | The behavior of the meerkat cannot be determined but the location of the individual is known. |
| Animal Not Visible | The meerkat is completely not visible and their location is unknown. |

### Data analysis

For each meerkat, the percent of visible time engaged in each behavior, height use, and posture was calculated by dividing the number of times each behavior was observed within an observation session by the total number of visible intervals within that session and multiplying by 100). Statistical analyses were performed in R version 4.6.1 (R Core Team 2026). A mean percent of time was calculated for the baseline and post-modification phases. These data were plotted using stacked column charts to visually illustrate differences between conditions.

Bayesian mixed-effects models were created to assess changes in sentry behavior, height use, and posture between the baseline and post-modification phases using the brms R package (Bürkner 2017). Phase was included as a fixed effect in each model, with baseline specified as the reference category. Meerkat ID and observation session were included as random intercepts in all models to account for non-independence among observations. Sentry behavior (0 = not observed, 1 = observed) and posture (0 = bipedal, 1 = tri/quadrupedal) were analyzed using generalized linear mixed-effects models with Bernoulli distributions and logit links. Height use was analyzed using a cumulative ordinal mixed-effects model, with height treated as an ordered categorical response (ground < low < high).

To assess whether crowd size was associated with sentry behavior, we fitted an additional Bayesian generalized linear mixed-effects model with a Bernoulli distribution and logit link. Crowd size was included as a continuous predictor and phase as a fixed effect to account for differences between study phases. Meerkat ID and observation session were included as random intercepts.

This study was approved by the Lincoln Park Research Committee, the governing body that oversees all research conducted at the institution.

## Results

There were 787 intervals collected on the meerkats across 90 observations during the baseline phase (Alanis: 151, Queen: 191, Miranda: 217, Sonny: 228). During the post-modification phase, there were 1,009 intervals collected on the meerkats (Alanis: 245, Queen: 266, Miranda: 236, Sonny: 262) across 113 observations.

Sentry behavior was more likely during the post-modification phase than during baseline (β = 2.90, 95% CrI [1.37, 4.49]; OR = 18.23, 95% CrI [3.94, 89.41]; P(β > 0) = 0.99). Crowd size was not clearly associated with the likelihood of sentry behavior. After accounting for study phase and repeated observations of individual meerkats, the estimated effect of crowd size on sentry behavior was small (β = 0.04, 95% CrI [−0.13, 0.22]; OR = 1.05, 95% CrI [0.88, 1.25]; P(β > 0) = 0.71). At the individual level, only Miranda was observed doing sentry duty (22% of visible observations; Figure 2) during the baseline phase. Following the addition of modifications to the meerkat enclosure, the other three meerkats were observed doing sentry duty (Alanis: 7% of visible observations, Queen: 20%, Sonny 40%; Figure 2); whereas Miranda decreased sentry behavior during this phase (8% decrease; Figure 2).

**Figure 2.**
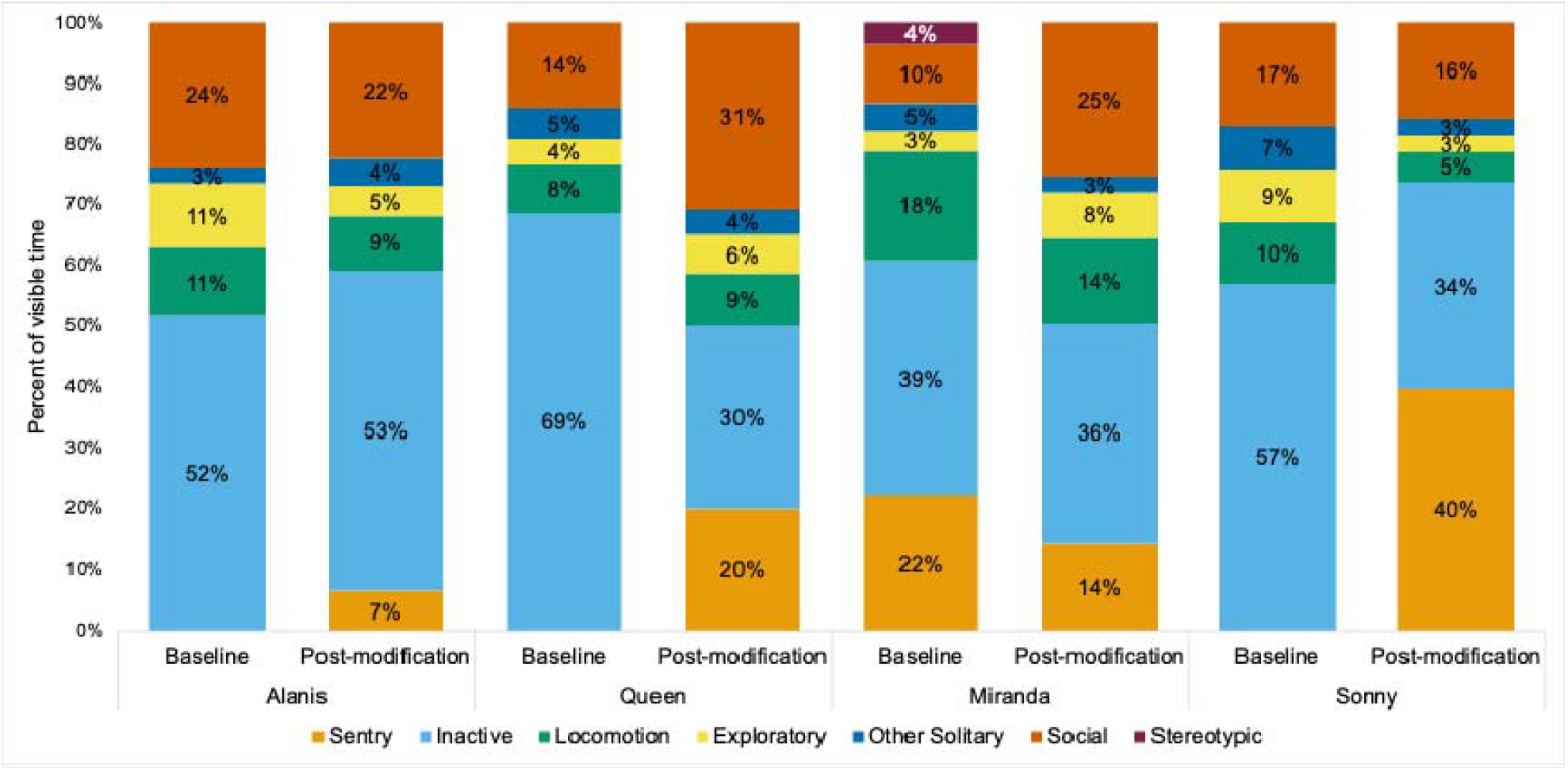
Simplified meerkat activity budget comparing baseline and post-modification phases. Inactive includes the behaviors of sitting, standing, lying, and leaning. Locomotion includes the behaviors of climbing, walking, and running. Exploratory includes the behaviors of digging, exploratory behavior, object manipulation, and play. Other solitary includes the behaviors of scent marking, vocalization, self-maintenance, feeding/foraging/drinking, and keeper interaction. Social includes the behaviors of social grooming, reproductive behavior, and huddling. For a detailed summary of individual behaviors (mean ± SD), see Supplementary Material 1.

The meerkats exhibited greater overall use of low-elevated locations during the post-modification phase compared with baseline. During baseline, 68.7% of visible observations occurred on the ground, 24.6% at low elevations, and 6.8% at high elevations, compared with 63.5%, 33.1%, and 3.4%, respectively, during the post-modification phase. The model provided some evidence that height use increased during the post-modification phase, but there was substantial uncertainty in the estimated effect (β = 0.39, 95% CrI [-0.46, 1.22]; OR = 1.48, 95% CrI [0.63, 3.38]; P(β > 0) = 0.82). At the individual level, Alanis and Sonny increased usage of low-elevated areas post-modification (26% and 29% increase, respectively; Figure 3), whereas Queen and Miranda decreased usage of elevated areas post-modification (8% and 20% decrease, respectively; Figure 3).

**Figure 3.**
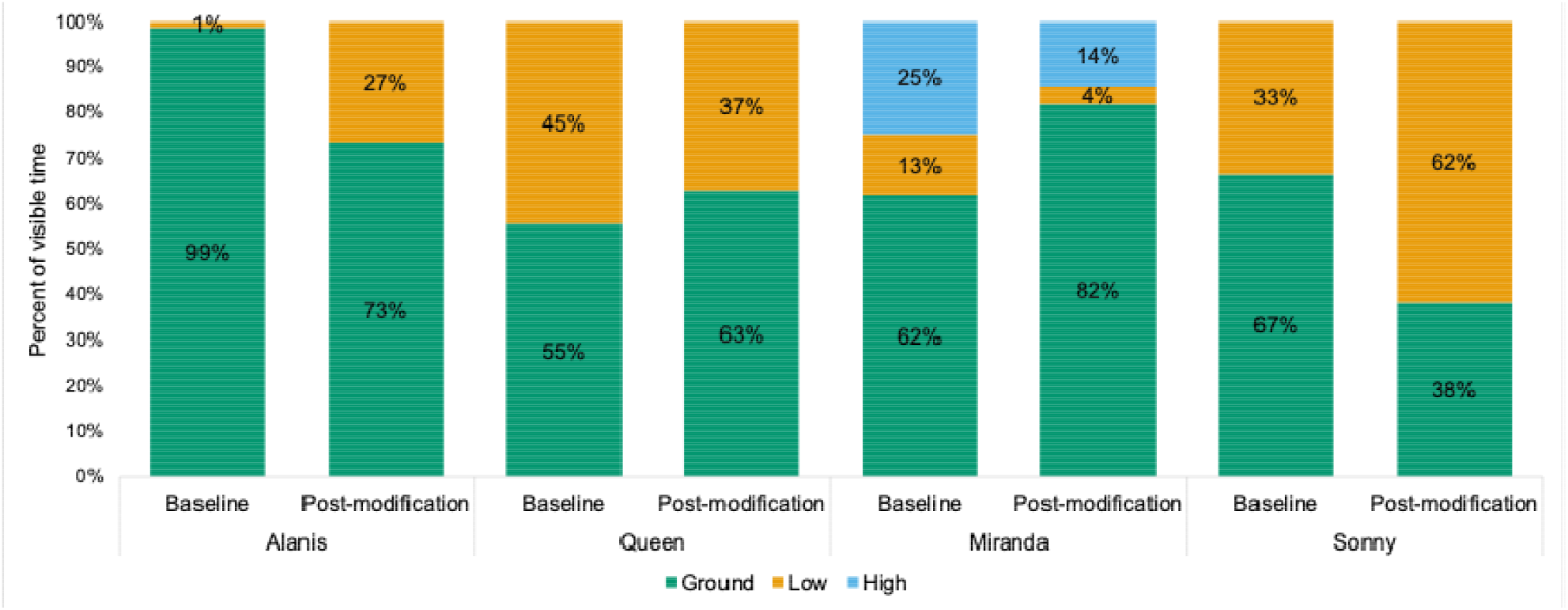
Meerkat height use comparing baseline and post-modification phases.

Posture differed between the baseline and post-modification phases. Tri/quadrupedal posture was observed in 33.3% of baseline observations and 51.0% of post-modification observations. The model indicated that tri/quadrupedal posture was more likely during the post-modification phase than during baseline, although there was uncertainty in the magnitude of the effect (β = 0.90, 95% CrI [-0.20, 1.99]; OR = 2.46, 95% CrI [0.82, 7.30]; P(β > 0) = 0.95). At the individual level, Alanis increased use of her tripedal posture after modifications were in place (32% increase; Figure 4). The other three meerkats decreased use of the bipedal posture in favor of the quadrupedal posture following modifications (Figure 4).

**Figure 4.**
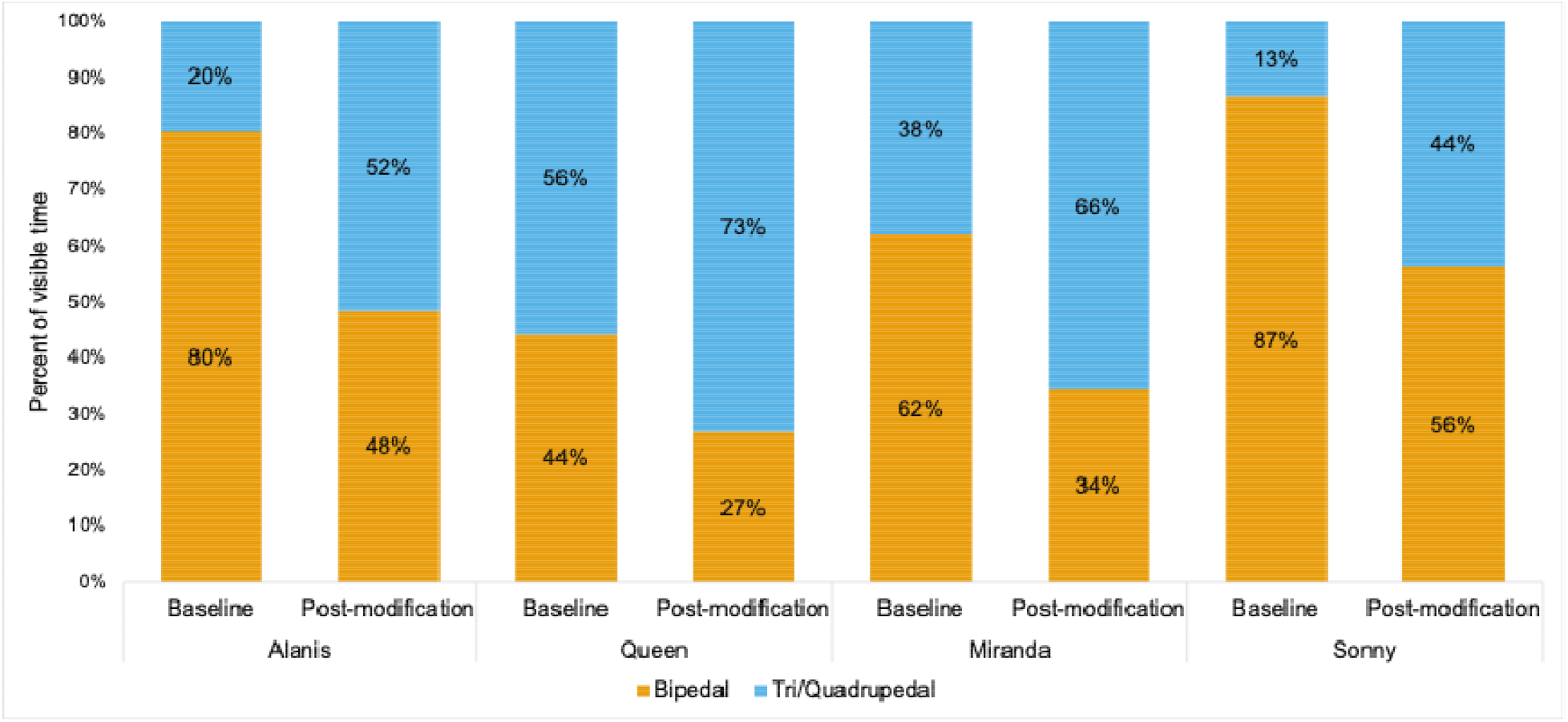
Meerkat posture when sitting or standing comparing baseline and post-modification phases.

## Discussion

Accessibility-focused modifications were associated with measurable changes in behavior and space use in this meerkat mob, which included individuals of advanced age as well as one individual with a disability. Sentry behavior, which typically involves climbing to elevated areas, increased for three of the four meerkats following the modifications. During the baseline phase, Miranda (female, 9 years old) was the only individual observed engaging in sentry behavior. However, following the modifications, the other three meerkats began participating in sentry duty, while Miranda’s sentry behavior decreased. In both zoo-housed and free-ranging meerkat mobs, some individuals engage in sentry behavior more frequently than others and may be referred to as “super sentinels” or “super guards” (Huels and Stoeger 2022; Rauber and Manser 2018; Tatalovic 2008). Nevertheless, individuals within a mob typically alternate as guards (Clutton-Brock et al. 1999; Manser 1999). Thus, the apparent absence of sentry rotation during the baseline phase is notable. The increased participation of the rest of the mob following the modifications suggests that previously inaccessible elevated areas may have limited opportunities for some members of the mob to engage in sentry behavior. Notably, the increase in sentry behavior for Alanis (female, 9 years old, limb amputation) following the modifications suggests that physical barriers within the enclosure, rather than her disability alone, may have previously limited opportunities to engage in this species-typical behavior. This finding highlights the importance of considering how enclosure design interacts with individual physical abilities to influence behavioral opportunities. By facilitating greater participation in sentry duty from the full mob, the modifications may have also reduced the concentration of sentry responsibilities on Miranda, allowing her more opportunities to engage in other behaviors, such as exploration and socialization (Figure 2). These findings illustrate how accessibility-focused enclosure modifications may influence not only individual behavior and space use but also the distribution of shared roles within a social group. Sentry behavior was not associated with crowd size in this mob, consistent with findings reported in other zoo-housed meerkats (Sherwen et al. 2014).

Following the modifications, Alanis and Sonny (male, 11 years old) increased their use of low-elevated areas, whereas Queen (female, 13 years old) and Miranda spent more time on the ground. These contrasting responses suggest that the modifications may have facilitated different patterns of enclosure use according to individual physical abilities and behavioral preferences. For Alanis, the increased use of elevated areas suggests that the modifications improved access that previously may have been difficult to navigate due to her limb amputation. Similarly, Sonny’s increased use of elevated areas suggests that the modifications may have expanded opportunities for him to access more areas within the enclosure. In contrast, Queen’s increased ground use may reflect age-related differences in mobility or preferences for particular enclosure areas. Miranda’s increased time on the ground may be related to her reduced participation in sentry duty and corresponding apparent increases in exploration and socialization. Together, these findings highlight the importance of considering individual differences when designing accessible environments, as modifications intended to improve access may not be equally effective or equally used by all individuals. For animals with disabilities, the effectiveness of enclosure modifications will likely depend on an individual’s physical abilities, willingness to use the modifications, and preferences for different areas of their enclosure.

Alanis increased her use of tripedal posture following modifications compared to the baseline phase. To our knowledge, no studies have examined behavioral or locomotor adaptation following limb amputation in meerkats or other members of Herpestidae. However, studies of domestic cats and dogs indicate that individuals can rapidly adapt their gait and weight bearing following limb amputation (Kamura et al. 2025; Galindo-Zamora et al. 2016). Importantly, baseline data collection began more than one year after Alanis’ amputation, providing substantial time for her to develop compensatory locomotor strategies and adjust to tripedal movement prior to this study. Therefore, the subsequent increase in tripedal posture following the modifications is unlikely to reflect adaptation to limb loss alone. Instead, Alanis’ increased use of tripedal posture following modifications suggests that changes to the enclosure may have facilitated greater use of this posture by enabling her to more readily navigate and interact with the environment. The other three meerkats increased their use of quadrupedal postures and decreased their use of bipedal postures following modifications, potentially reflecting age-related changes in balance or stability. However, these changes may also be related to differences in behavioral activities or space use following the modifications. For example, the increase in quadrupedal posture may have been the result of a reduced need to maintain vigilance from the ground due to the addition of low-elevation perches. Modifications are likely to impact how individuals physically interact with their environment and may facilitate the adoption of different postures as they navigate available spaces.

Collectively, the findings of this study indicate that an animal’s physical condition alone does not determine whether they can perform a species-typical behavior. Rather, the accessibility of their environment may determine whether an animal’s physical condition becomes behaviorally limiting. This is parallel to the person-environment fit model of disability in humans, which emphasizes that mismatches between a person’s physical condition and their environment create a disability, as opposed to the physical condition itself (Webb et al. 2011). For the meerkats of this study, the modifications provided intermediate height opportunities, which allowed individuals to obtain some benefits of elevation while requiring less demanding climbing effort than the existing termite mound. Universal design and accessibility-focused retrofitting is intended to provide multiple usable options for individuals of diverse abilities (Story 2001), not produce identical use of or behavioral responses to the physical space. The modifications provided the meerkats with more opportunities to elevate themselves, but they were not used equivalently by all individuals. Thus, effectiveness should not necessarily be judged by whether every individual increases use of a particular structure, but rather if it provides accessible opportunities for individuals to engage with their environment. In doing so, animals may be provided with more choice and control in how they interact with their environment, which helps promote positive welfare (Englund and Cronin 2023).

Zoo environments have traditionally been designed around species-level needs, yet individuals within a species can vary substantially in their physical and sensory abilities, influencing how effectively they can access and interact with their environment. Aging, injury, and other changes in physical or sensory function can alter an animal’s ability to navigate an enclosure that may otherwise be considered appropriate for the species. Consequently, accessibility should be considered not only as a response to an identified limitation, but as a proactive component of enclosure design that accommodates individual variation and changing abilities across an animal’s lifetime. Incorporating flexible design features, such as modular structures, low-gradient slopes, intermediate levels between elevated areas, and non-slip surfaces, may help maintain access to important resources and opportunities for species-typical behavior as animals’ abilities change.

There are a few limitations to acknowledge in this study. First, this study examined a single group of four meerkats, including only one individual with a limb amputation, limiting the generalizability of the findings to other meerkats, types of disabilities, species, or facilities. Second, the pre/post study design cannot fully exclude the influence of temporal changes unrelated to the enclosure modifications, as baseline and post-modification observations were conducted one year apart. Future research should build on these findings through multi-institutional studies involving larger numbers of animals with diverse disabilities and accessibility needs. Preference or choice-based assessments could help determine which accessible structures animals choose to use, while measures of locomotion, gait, balance, and physical effort could provide additional insight into how modifications affect the physical demands of navigating an enclosure. Longer-term monitoring is also needed to determine whether accessible features continue to be used and provide benefits over time.

This study demonstrates that relatively simple enclosure modifications can meaningfully change how animals interact with their environment. Providing intermediate-height structures was associated with changes in sentry behavior, height use, and posture, although responses varied among individuals. Rather than viewing disability solely as a characteristic of an individual animal, these findings highlight the importance of considering how an animal’s abilities interact with their environment. Designing for a range of abilities may help maintain access to important resources and opportunities for species-typical behavior. Although the modifications in this study were implemented to improve accessibility for individuals with physical limitations, the observed behavioral changes across the mob suggest that accessible design may benefit other group members as well. Accessibility-focused enclosure design represents a promising and practical component of evidence-based care that can enhance animal welfare.

## Acknowledgements

Thank you to Alex Kim, Isabel DePaz, Isabella Robinson, and Audrey Rumney for supporting data collection. Thank you to Cassy Kutilek, Rachel Valentas, Maria Sheehan, Chris Garcia, Anthony Nielsen, Marisa Elizalde, Raven Spencer, Bridget Czarnik, Amanda Kalinowski, Rebecca Woolley, Arianna Ewing, Lupe Rojas, and Dan Kozlowski for caring for the meerkats and supporting modifications in the meerkat enclosure. Thank you to the facilities staff who prepared the modifications and to Jason Wark, Tasha Wierzal, Maureen Leahy, and Dave Bernier for supporting this research. Thank you to Bill Green for continued discussions related to animals with disabilities.

