## Supplementary Table 1 for "Assessing Enclosure Modifications to Enhance Accessibility for Zoo-Housed Meerkats (*Suricata suricatta*)"

**Supplementary Table 1.** Percentage of visible time spent in each behavior by individual meerkats during baseline and post-modification phases. Values are mean ± SD (%) across visible observations.

| **Behavior** | **Alanis** | | **Queen** | | **Miranda** | | **Sonny** | |
| --- | --- | --- | --- | --- | --- | --- | --- | --- |
|  | **Baseline** | **Post** | **Baseline** | **Post** | **Baseline** | **Post** | **Baseline** | **Post** |
| Climbing | 0.00 ± 0.00 | 0.00 ± 0.00 | 0.80 ± 2.77 | 0.69 ± 3.71 | 0.98 ± 3.26 | 0.00 ± 0.00 | 0.46 ± 2.27 | 0.36 ± 1.89 |
| Digging | 2.36 ± 5.72 | 1.11 ± 3.27 | 2.89 ± 8.77 | 1.42 ± 4.49 | 0.87 ± 2.88 | 1.47 ± 3.67 | 5.52 ± 14.48 | 0.00 ± 0.00 |
| Exploratory Behavior | 8.67 ± 9.53 | 2.54 ± 5.25 | 0.80 ± 2.77 | 2.87 ± 6.20 | 1.85 ± 4.14 | 3.56 ± 7.16 | 3.17 ± 5.08 | 1.79 ± 4.76 |
| Feeding/ Foraging/ Drinking | 1.90 ± 4.46 | 0.36 ± 1.89 | 0.00 ± 0.00 | 2.76 ± 9.96 | 1.74 ± 8.34 | 1.43 ± 5.91 | 1.25 ± 6.12 | 0.00 ± 0.00 |
| Huddling | 23.47 ± 25.98 | 20.16 ± 34.27 | 10.93 ± 23.56 | 25.69 ± 35.30 | 7.89 ± 16.47 | 18.61 ± 32.50 | 14.05 ± 25.17 | 10.04 ± 25.39 |
| Keeper Interaction | 0.00 ± 0.00 | 1.43 ± 7.56 | 0.00 ± 0.00 | 1.38 ± 7.43 | 0.00 ± 0.00 | 0.00 ± 0.00 | 0.00 ± 0.00 | 0.00 ± 0.00 |
| Leaning | 0.00 ± 0.00 | 10.22 ± 20.51 | 0.00 ± 0.00 | 0.00 ± 0.00 | 0.00 ± 0.00 | 0.00 ± 0.00 | 0.00 ± 0.00 | 0.36 ± 1.89 |
| Lying | 9.81 ± 17.78 | 16.15 ± 25.75 | 9.53 ± 21.75 | 8.39 ± 15.08 | 4.40 ± 8.47 | 1.43 ± 4.48 | 7.85 ± 21.53 | 3.57 ± 9.11 |
| Object Manipulation | 0.00 ± 0.00 | 0.00 ± 0.00 | 0.00 ± 0.00 | 0.43 ± 2.32 | 0.00 ± 0.00 | 0.00 ± 0.00 | 0.00 ± 0.00 | 0.00 ± 0.00 |
| Pacing | 0.00 ± 0.00 | 0.00 ± 0.00 | 0.00 ± 0.00 | 0.00 ± 0.00 | 3.48 ± 11.12 | 0.00 ± 0.00 | 0.00 ± 0.00 | 0.00 ± 0.00 |
| Play | 0.00 ± 0.00 | 0.75 ± 2.77 | 0.00 ± 0.00 | 1.47 ± 5.96 | 0.48 ± 2.32 | 1.79 ± 4.76 | 0.42 ± 2.04 | 0.75 ± 2.77 |
| Reproductive Behavior | 0.00 ± 0.00 | 0.00 ± 0.00 | 0.00 ± 0.00 | 0.38 ± 2.06 | 0.00 ± 0.00 | 0.00 ± 0.00 | 0.00 ± 0.00 | 0.00 ± 0.00 |
| Running | 1.73 ± 3.98 | 0.00 ± 0.00 | 0.00 ± 0.00 | 0.57 ± 3.09 | 3.04 ± 8.76 | 0.36 ± 1.89 | 0.00 ± 0.00 | 0.45 ± 2.36 |
| Scent marking | 0.00 ± 0.00 | 0.00 ± 0.00 | 0.00 ± 0.00 | 0.00 ± 0.00 | 0.43 ± 2.09 | 0.00 ± 0.00 | 1.39 ± 3.75 | 0.75 ± 2.77 |
| Self-grooming | 0.79 ± 3.37 | 0.00 ± 0.00 | 1.24 ± 4.49 | 0.00 ± 0.00 | 2.27 ± 5.34 | 0.00 ± 0.00 | 3.95 ± 7.88 | 0.00 ± 0.00 |
| Self-maintenance/ Comfort Behaviors | 0.00 ± 0.00 | 2.30 ± 6.90 | 0.00 ± 0.00 | 1.03 ± 4.09 | 0.00 ± 0.00 | 1.55 ± 6.51 | 0.00 ± 0.00 | 2.02 ± 6.04 |
| Sentry | 0.00 ± 0.00 | 5.79 ± 15.28 | 0.00 ± 0.00 | 19.20 ± 34.46 | 22.74 ± 31.88 | 14.08 ± 30.09 | 0.00 ± 0.00 | 41.25 ± 37.41 |
| Sitting | 22.99 ± 29.60 | 21.84 ± 28.99 | 50.32 ± 29.86 | 22.61 ± 25.88 | 26.31 ± 24.61 | 26.77 ± 27.09 | 27.81 ± 30.09 | 22.57 ± 27.42 |
| Social Grooming | 0.56 ± 2.36 | 0.40 ± 2.10 | 0.00 ± 0.00 | 2.41 ± 11.23 | 2.27 ± 5.34 | 2.86 ± 6.00 | 2.75 ± 7.98 | 5.08 ± 18.98 |
| Standing | 14.81 ± 22.50 | 8.43 ± 14.17 | 9.50 ± 16.30 | 0.57 ± 3.09 | 7.44 ± 10.98 | 13.49 ± 26.89 | 21.22 ± 23.93 | 7.01 ± 12.73 |
| Vocalization | 0.00 ± 0.00 | 0.00 ± 0.00 | 3.34 ± 8.66 | 0.00 ± 0.00 | 0.00 ± 0.00 | 0.00 ± 0.00 | 0.42 ± 2.04 | 0.00 ± 0.00 |
| Walking | 12.90 ± 24.49 | 8.52 ± 11.52 | 10.63 ± 17.33 | 8.12 ± 10.93 | 13.80 ± 15.29 | 12.61 ± 16.52 | 9.75 ± 10.47 | 4.01 ± 9.71 |
